# Single-Cell Transcriptional and Epigenetic Profiles of Male Breast Cancer Nominate Salient Cancer-Specific Enhancers

**DOI:** 10.1101/2022.12.02.518918

**Authors:** Hyunsoo Kim, Kamila Wisniewska, Matthew J. Regner, Philip M. Spanheimer, Hector L. Franco

## Abstract

Male breast cancer represents about 1% of all breast cancer diagnoses and, although there are some similarities between male and female breast cancer, the paucity of data available on male breast cancer makes it difficult to establish targeted therapies. To date, most male breast cancers (MBC) are treated according to protocols established for female breast cancer (FBC). Thus, defining the transcriptional and epigenetic landscape of MBC with improved resolution is critical for developing better avenues for therapeutic intervention. In this study, we present matched transcriptional (scRNA-seq) and epigenetic (scATAC-seq) profiles at single-cell resolution of two treatment naïve MBC tumors processed immediately after surgical resection. These data enable the detection of differentially expressed genes between male and female breast tumors across immune, stromal, and malignant cell types, to highlight several genes that may have therapeutic implications. Notably, *MYC* target genes and *mTORC1* signaling genes were significantly upregulated in the malignant cells of MBC compared to the female counterparts. To understand how the regulatory landscape of MBC give rise to these male-specific gene expression patterns, we leveraged the scATAC-seq data to systematically link changes in chromatin accessibility to changes in gene expression within each cell type. We observed cancer-specific rewiring of several salient enhancers and posit that these enhancers have a higher regulatory load than lineage specific enhancers. We highlight two examples of previously unannotated cancer-cell specific enhancers of *ANXA2* and *PRDX4* gene expression and show evidence for super-enhancer regulation of *LAMB3* and *CD47* in male breast cancer cells. Overall, this dataset annotates clinically relevant regulatory networks in male breast tumors, providing a useful resource that expands our current understanding of the gene expression programs that underlie the biology of MBC.

## INTRODUCTION

Male breast cancer (MBC) is a rare type of cancer that occurs in the breast tissue of men. MBC accounts for only 1% of total breast cancer incidence^1,2^, however, men are likely to present with larger, higher-grade tumors, and more lymph node involvement compared to females with breast cancer^3^. Moreover, the 5-year mortality of MBC is higher than that of female breast cancer (FBC)^4^, which may be due to the older age at the time of diagnoses, delays in diagnosis, the presence comorbidities, or intrinsic biological differences^3-5^. Most men with breast cancer are diagnosed with invasive ductal carcinoma, and their tumors are estrogen-receptor (ER) positive, progesterone-receptor (PR) positive, and HER2 negative^5-8^. At the molecular level, most men will present with Luminal A-like or Luminal B-like tumors^6,9^, with some studies suggesting that MBCs have unique subtypes, M1 and M2, that differ from the intrinsic subtypes of FBC^10,11^. As with FBC, MBC risk increases for men with familial history of *BRCA* mutations. However, men show higher risk for *BRCA2* mutations than *BRCA1* mutations compared to women^12,13^. Men are also at higher risk of developing breast cancer if they are African American^14^ or have comorbidities such as Klinefelter’s syndrome, hormone imbalances, liver disease, and obesity, among others^5,15,16^.

Although male breast cancer presents with similarities to certain female breast cancers, the paucity of data related to the treatment of males makes it difficult to find targeted therapies^17-19^. Instead, men with breast cancer are treated in accordance with treatment paradigms for women, even if the efficacy of these treatments is low^18,20^. Interestingly, studies comparing the genomic profiles of male and female breast cancers have found important differences that are potentially driving outcomes of their respective disease^10,11,21,22^. For example, several microarray-based studies have found that *NAT1*^10^, *mTOR*^23^, *EIF4E*^23^, *THY1*^11^, and *SPAG5*^11^ are upregulated in MBC compared to FBC and may serve as prognostic biomarkers. These initial studies highlight the fundamental differences of male and female breast cancer and provide the impetus for defining male-specific disease mechanisms that will lead to better treatment options.

Single-cell genomics have revolutionized our ability to investigate the cellular, transcriptional, and epigenetic heterogeneity of human tumors with improved resolution. Single-cell RNA-seq (scRNA-seq)^24-29^ refines our ability to measure the transcriptional profiles of thousands of individual cells within a particular tumor specimen to make conclusions about the underlying cellular heterogeneity of tumors and pinpoint salient gene expression programs. These gene expression programs are controlled and sustained by regulatory elements (e.g., *cis*-acting enhancer elements) scattered throughout the genome that are often rewired and repurposed by cancer cells to drive oncogenic transcriptional programs^30-33^. The chromatin accessibility landscape may now be robustly profiled thanks to recent improvements in single-cell sequencing assay for transposase-accessible chromatin (scATAC-seq), revealing several layers of gene regulation, including cisregulatory elements^34,35^. Together with scRNA-seq, scATAC-seq offers unprecedented resolution to reveal novel gene regulatory mechanisms in MBC. Although there are some notable cancer datasets with matched scRNA-seq and scATAC-seq^36,37^, none have been reported for human MBC. Herein we present matched transcriptional (scRNA-seq) and epigenetic (scATAC-seq) profiles of two treatment naïve MBC tumors processed immediately after surgical resection. First, we defined the differentially expressed genes between male and female breast tumors, specifically within the malignant cell types, to highlight several genes that may have therapeutic implications. Then, we systematically linked changes in chromatin accessibility to changes in gene expression to annotate the enhancer landscape within each cell type. Finally, we highlight the cancer-specific rewiring of several salient enhancers that drive abnormally high levels of male-specific gene expression and investigate the transcription factor occupancy at enhancers. Together these data enable the annotation of the cellular composition, transcriptional, and epigenetic landscape of male breast tumors to help pinpoint drivers of this rare disease.

## RESULTS

### Matched scRNA-seq and scATAC-seq of male breast cancer

Two treatment-naïve male breast cancer patients underwent mastectomy with curative intent (Fig. 1). Immediately following surgical resection, each tumor was gently dissociated into a suspension of live cells using a gentle collagenase and hyaluronidase digestion and prepped for lipid droplet-based scRNA-seq and scATAC-seq via the 10x Genomics Chromium system (Fig. 1A). Each tumor specimen was divided into two pools to generate independent scRNA-seq and scATAC-seq libraires. Since these tumor specimens were never frozen or fixed in any way, a high level of cell viability during the dissociation process was maintained for robust sequencing coverage in single cells.

**Figure 1.**
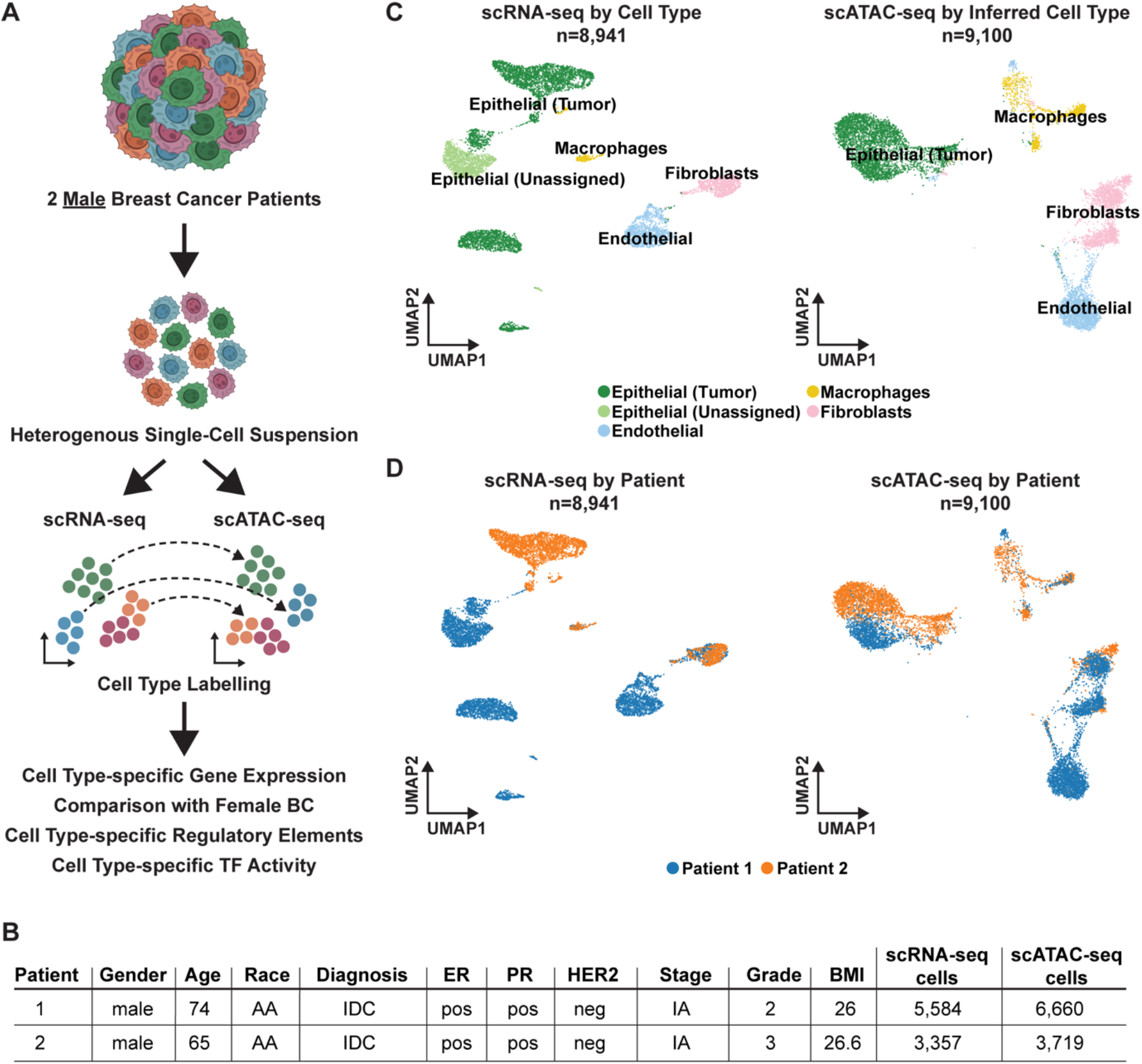
Overview of scRNA-seq and scATAC-seq of Male Breast Cancer Tumors. A. Cartoon of scRNA-seq and scATAC-seq workflow. B. Table capturing clinical features of the male breast tumors and total cells captured for both scRNA-seq and scATAC-seq. Abbreviations; AA = African American, IDC = Infiltrating Ductal Carcinoma, pos = positive immunohistochemical staining, neg = negative for immunohistochemical staining. C. UMAP plot of scRNA-seq colored by cell type (*left*) and scATAC-seq colored by inferred cell type (*right*) across both male breast cancer patients capturing total cells after QC (*see methods*). D. UMAP plot of scRNA-seq (*left*) and scATAC-seq (*right)* colored by patient of origin.

After quality control and doublet removal for each patient dataset, we obtained 8,941 total cells profiled by scRNA-seq (Supplementary Fig. S1A) and 10,379 total cells profiled by scATAC-seq (Supplementary Fig. S1E). To ensure the robustness of our downstream analysis, we removed any cell type clusters that had less than 5,000 RNA or ATAC fragment counts on average, or could not confidently be assigned a cell type label (s*ee Methods*; Supplementary Fig. S1C and S1G). After applying these filters, we proceeded with a dataset composed of 8,941 profiled by scRNA-seq cells and 9,100 total cells profiled by scATAC-seq (Fig. 1B).

To annotate the transcriptional profiles of the various cell types within these MBC tumor specimens, we performed principal component analysis (PCA) using the top 2,000 most variable genes across all 8,941 scRNA-seq cells. To correct for any technical variation, *Harmony* batch correction^38^ was applied to the dataset and the cells were then classified into transcriptionally distinct clusters with graph-based clustering using the top 30 PCs and visualized using a Uniform Manifold Approximation and Projection (UMAP) plot. We identified four major cell types (epithelial cells, macrophages, fibroblasts, and endothelial cells) (Fig. 1B, *left*) across nine clusters (Suppl. Fig. S1A). Notably, the non-malignant cell types, such as the macrophages and fibroblasts, showed intermixing of cells from both patients, suggesting that these cell types had similar transcriptional profiles across patients. Conversely, the epithelial cell types (which were later confirmed to be malignant via inferCNV) showed patient-specific transcriptional profiles (Fig. 1C, *left*). Because these data were batch corrected prior to clustering, we believe that the patient-specific transcriptional profiles of the epithelial cell types are biologically meaningful.

To analyze the chromatin accessibility landscape (*vis-a-vis* the epigenetic landscape) of these tumors, the scATAC-seq data was processed by creating a matrix of contiguous genomic tiles across the genome, in which we quantified Tn5 insertion counts across every cell. Then we performed iterative latent semantic indexing (LSI) on the top 25,000 most variable genomic tiles ^37,39^. We used Seurat v4.0.5 cross-modality integration^40^ with the top 30 LSI dimensions (constrained to cells of the same patient tumor) to assign cell type labels from the matching scRNA-seq data to the scATAC-seq cells, and visualized the cells in the UMAP plot 4 (Fig. 1C, *right*)^41^. The scATAC-seq cells were clustered mostly by cell type and not by patient, which showed the quality of the dataset and data processing pipeline (Fig. 1D, *right*). All four cell types (epithelial cells, macrophages, fibroblasts, and endothelial cells) were also observed in scATAC-seq (Fig. 1C, *right*). Similar to the scRNA-seq, we found that the cell type clusters of macrophages and fibroblasts contained cells from both patients, while the cell type clusters of epithelial cells were patient-specific in scATAC-seq (Fig. 1D, *right*). These observations likely reflect the biological overlap of the non-malignant cells across all patients and highlights the unique, and possibly tractable, biological features of the malignant cells within each patient’s tumor.

### Transcriptional profiles of male versus female ER+ malignant epithelial cells

To determine the transcriptional differences between male and female cancer cells, we first identified the malignant epithelial cell types within each patient by inferring putative copy number events for each cell cluster using the inferCNV^42^ approach (Supplementary Fig. 2A-C). Then we compared the cancer-epithelial cell profiles of our MBC tumors to publicly available scRNA-seq profiles of FBC epithelial cells^43^ of the same molecular subtype. The male epithelial cells from both patients grouped together in a UMAP that contained both male and female cancer-epithelial cells, suggesting they have distinct transcriptional profiles compared to the female epithelial cells (Fig. 2A). Differential gene expression analysis between MBC and FBC, at a single-cell resolution, identified 1,004 upregulated genes and 14 downregulated genes in MBC with log2FC > 0.25 and adjusted P-value < 0.01 (Supplementary Table S1). To simplify the heatmap visualization, we filtered the differentially expressed genes with an adjusted P-value < 1e-12 to arrive at the top 25 upregulated genes and top 5 downregulated genes (Fig. 2B). To validate the soundness of the differential gene expression analysis, we visualized the expression of the top two upregulated genes (*RPS4Y1* and *TNFRSF12A*) and top two downregulated genes (*FOS* and *XIST*) in the UMAP plot (Fig. 2C). Sex-specific genes such as *RPS4Y1* (located on the Y-chromosome) and XIST (needed for X-chromosome inactivation^44^) were exclusively expressed in the corresponding male and female epithelial cells, giving confidence to our analysis (Fig. 2C). Gene ontology analysis^45^ of the differentially expressed genes showed that the upregulated genes in MBC are enriched in ‘*MYC targets*’ and ‘*mTORC1 signaling*’ which suggests the possibility of using BET bromodomain inhibitors^46^ and/or mTOR inhibitors^47,48^ for MBC treatment (Fig. 2D, Supplementary Table S1). Conversely, the downregulated genes were enriched in ‘*Estrogen Response Late*’ pathway, suggesting that although the male epithelial cells are ER+, the estrogen driven gene expression pathways may be muted. Together, these data reveal sex-specific gene expression differences specifically within the malignant cell types of male and female breast cancer.

**Figure 2.**
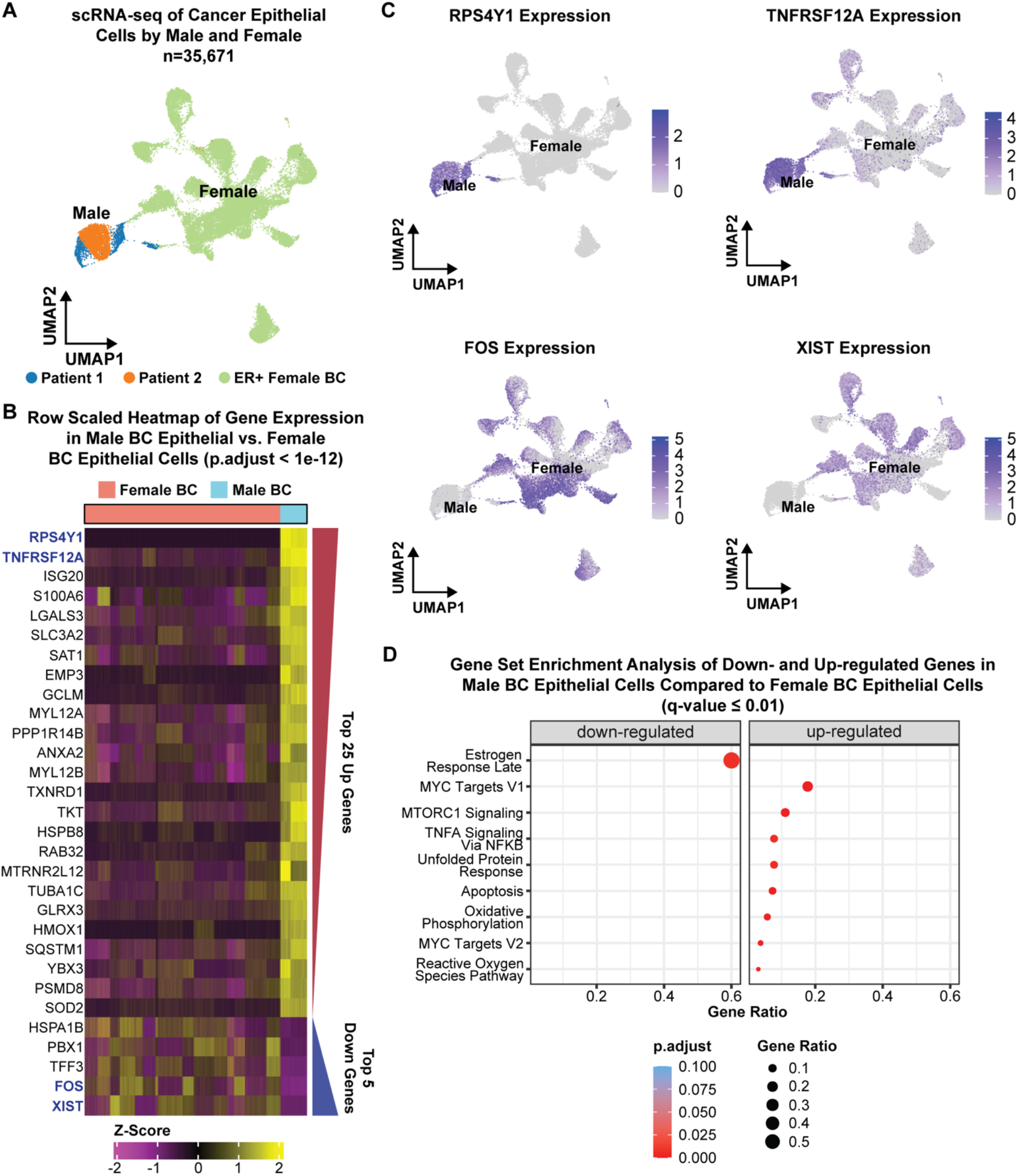
scRNA-seq Analysis Comparing Male and Female Breast Cancer. A. scRNA-seq UMAP plot of epithelial cells colored by male patients from this study and female patients from Pal *et al*. ^43^. B. Row scaled heatmap of gene expression in male cancer epithelial cells versus female cancer epithelial cells. The heatmap shows the top 25 upregulated genes and the top 5 downregulated genes with p-value adjusted < 1e-12. Red/blue wedges to the right of the heatmap correspond to the p-value adjusted trends among the upregulated and downregulated groups. More significant upregulated genes are at the top of the heatmap while the most significant downregulated genes are at the bottom of the heatmap. C. UMAP plots of scRNA-seq male and female cancer epithelial cells colored by normalized expression of the two most significantly upregulated genes in male breast cancer (*RPS4Y1* and *TNFRSF12A*) and the two most significantly downregulated genes in male breast cancer (*FOS* and *XIST*) compared to female breast cancer. D. Hallmark gene set enrichment analysis of downregulated and upregulated genes in male breast cancer tumors compared to female breast cancer tumors (q-value ≤ 0.01).

### Systematic annotation of cancer-specific enhancer elements in MBC

Chromatin accessibility is a prerequisite for gene transcription and for the activity of the regulatory elements that regulate transcription. Moreover, chromatin accessibility is cell-type specific, usually constrained to the gene expression patterns of a particular cell lineage. Thus, it is possible to deconvolute the *cis* regulatory elements that drive cell-type specific gene expression within MBC tumors using scATAC-seq. First, we called statistically significant chromatin accessibility peaks across all cells and found that the majority of peaks were located in intronic regions (46.2%) or distal intergenic regions (31.3%) (Fig. 3). Next, we linked changes in chromatin accessibility to changes in gene expression by performing a peak-to-gene correlation analysis (see Methods; Fig. 3)^39^. Briefly, we aggregated the sparse peak counts within groups of similar scATAC-seq cells (∼100 cells per group), identified via *k*-nearest neighbors, to generate more informative metacell observations for each peak in the analysis. Then we computed the correlation between the accessibility of every peak and the expression of every gene across scATAC-seq cells imputed after the Seurat v4.0.15 label transfer procedure^40^. Overall, the peak-to-gene linkage analysis identified 11,719 unique distal peaks participating in 22,869 distal peak-to-gene links in *cis* across all cell types (Pearson correlation ≥ 0.45 and FDR < 1e-12). We further categorized these peak-to-gene links into five *k*-means clusters before visualizing them in heatmap form where we observed highly consistent patterns between gene expression and linked peak accessibility (Fig. 3B). Furthermore, the *k*-means clustering of the peak-to-gene links revealed cell-type-specific enhancer-gene pairs that were characteristic of each cell type; macrophages (cluster 1), epithelial-tumor cells (clusters 2 and 3), fibroblasts (cluster 4), and endothelial cells (cluster 5) (Fig. 3B). Herein, we refer to these linked distal peaks as putative enhancers of their paired target genes (Supplementary Table S2).

**Figure 3.**
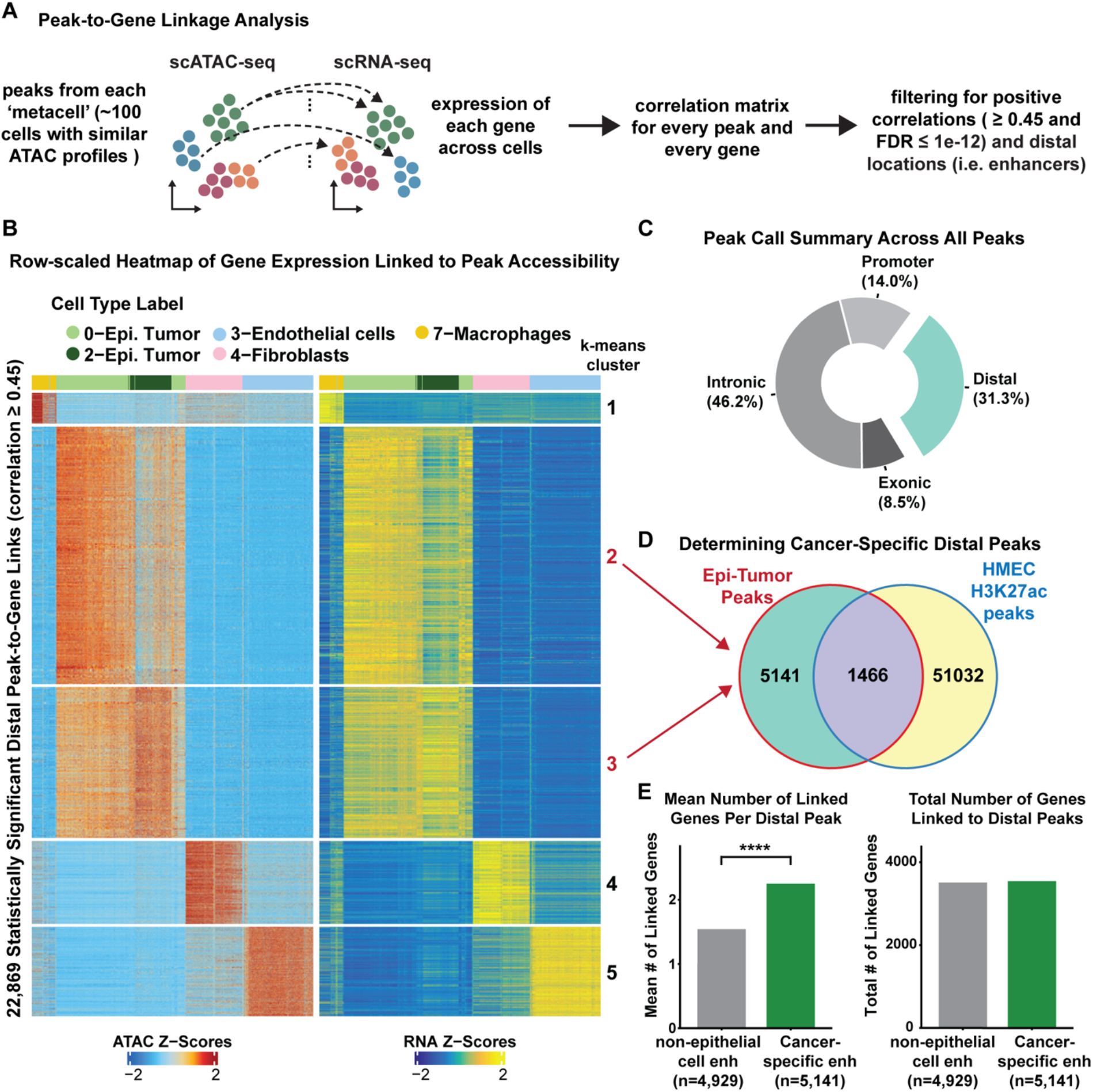
Identification of cancer-specific enhancers in male breast cancer. A. Workflow for the enhancer peak-to-gene identification strategy. B. Row-scaled heatmap of statistically significant distal peak-to-gene links (FDR<1e-12). Rows represent the accessibility of a distal peak and the expression of its linked gene. Columns represent cell types. Cancer-epithelial peak-to-gene links are clustered by k-means clustering and corresponding cluster numbers are denoted in red at the right side of the heatmap. C. Peak call summary pie chart showing the location of all peaks. D. Venn-diagram showing the number of cancer-specific distal peaks (i.e., 5,141 peaks in clusters 2 and 3 from Fig. 3B) that do not overlap with regulatory elements found in normal human mammary epithelial cells (HMEC). E. *Left*: Bar chart depicting the mean number of linked genes per distal peak in cancer-specific distal peaks (green, n=5,141) vs. all non-epithelial peaks (grey, n=4,929) cells. **** denote *p*-value < 2.22e-16 (Wilcoxon rank-sum test). *Right*: Bar chart depicting the total number of genes linked to cancer-specific distal peaks (3,586 total genes) vs non-epithelial distal peaks (3,551 total genes).

To differentiate between cancer-specific enhancers and lineage-specific enhancers, we extracted the genomic coordinates for each peak that was enriched in the cancer-epithelial cell specific k-means groups (clusters 2 and 3; Fig 3B) and overlapped these with previously annotated enhancer elements in normal human mammary epithelial cells (HMECs)^49^. We found a total of 5,141 cancer-specific enhancers (participating in 11,551 peak-to-gene links) that were not present in normal mammary epithelial cells and specifically active in the cancer-epithelial cells of MBC (Fig. 3D, Supplementary Table S2). Interestingly, the cancer-specific enhancers tend to link to more genes on average (∼2.2 genes per enhancer) compared to the non-cancer enhancers found in all other cell types (∼1.7 genes per enhancer) (Fig. 3E). This suggests that the cancer-specific enhancers may have a higher regulatory load, being associated to the expression of more genes, as compared to lineage-specific enhancers. We also note that transcription factor motif enrichment analysis of the cancer-specific enhancers revealed *FOXA1, MESP1/2*, and *TFAP2C* as the top three transcription factors enriched at these enhancers based on adjusted P-value (Supplementary Table S3). These data point to a significant rewiring of enhancer elements in cancer-epithelial cells compared to normal epithelial cells, that can potentially sustain oncogenic transcription in MBC.

### Tumor epithelial cells acquire enhancer elements that drive genes involved in cancer progression

The peak-to-gene analysis revealed a striking amount of cancer-specific enhancer-gene pairs that could potentially serve as biomarkers or even tractable pathways in male breast cancer. Therefore, we wanted to identify the cancer-specific enhancers that were linked to male breast cancer genes that are up-regulated in comparison to female breast cancer. In total, we found 61 genes that are upregulated in MBC compared to FBC, whose enhancers are specifically active in the cancer-epithelial cell fraction of MBC tumors (Fig. 4A and B). One of the highest expressed genes is *ANXA2*, a gene involved in tumor heterogeneity and cancer progression^50^. It has been reported that *ANXA2* is more expressed in African American triple negative breast cancer (TNBC) patients compared to Caucasian TNBC patients, and high expression of *ANXA* is associated with worse survival^51^. The enhancer linked to *ANXA2* is significantly enriched in the cancer-epithelial cells but not in the non-malignant cell types (log2FC = 2.3 and FDR < 0.001), resulting in significantly higher expression of *ANXA2* as measured by the scRNA-seq (Wilcoxon rank sum tests, p-value < 2.22e-16) (Fig. 4C). Of note, this enhancer is not normally active in human mammary epithelial cells (HMEC) and, perhaps more interestingly, is also not annotated in the ENCODE Consortium’s curated registry of cis-regulatory elements compiled across hundreds of different cell types^49^ (Fig. 4C). Furthermore, motif analysis revealed *YY1* as the most significantly enriched transcription factor at this enhancer and *MAFF* as the highest enriched transcription factor at the promoter of *ANXA2* (Fig. 4D). *YY1* is an important mediator of enhancer-promoter interactions^52^, and the expression of *YY1* and *MAFF* in the cancer-epithelial cells was confirmed using the matching scRNA-seq dataset (Fig. 4D). Taken together, this previously un-annotated cancer-specific enhancer represents a novel regulator of *ANXA2* in MBC.

**Figure 4.**
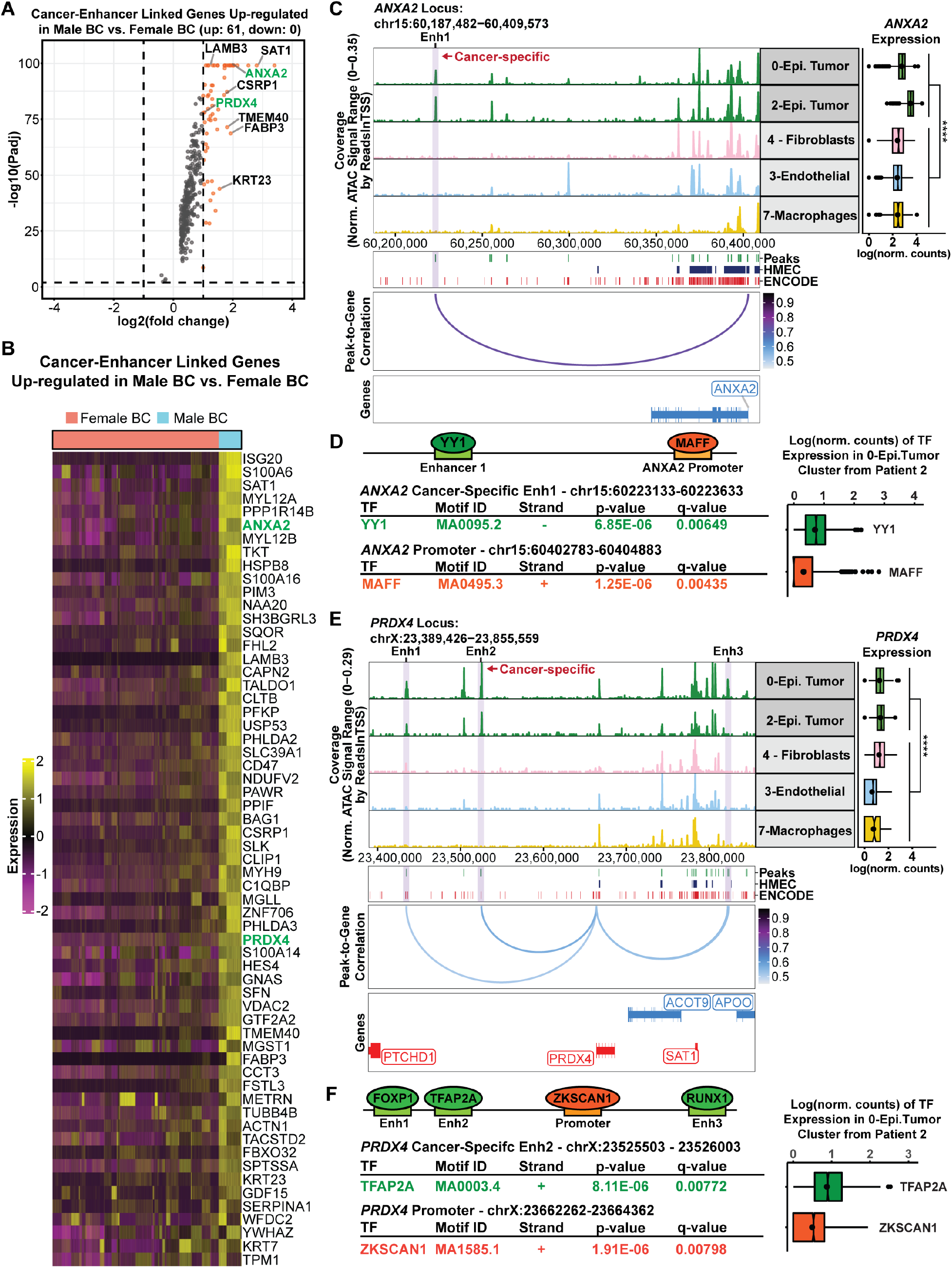
Cancer-specific enhancers drive the expression of *ANXA2* and *PRDX4 in male breast cancer*. A. Volcano plot of cancer-enhancers linked to genes that are upregulated in male breast cancer cells versus female breast cancer cells. B. Row-scaled heatmap of all 61 upregulated cancer-enhancer linked genes in male breast cancer versus female breast cancer. More significant upregulated genes are at the top of the heatmap. C. Browser track view of chromatin accessibility at the *ANXA2* locus across epithelial cancer cells (Epi. Tumor) and all other cell-type clusters. The cancer-specific enhancer 1 (Enh1) peak that is linked to *ANXA2* gene expression is highlighted in light purple. The peak track below the browser track denotes all scATAC-seq peaks from this study (Peaks), regulatory elements found in human mammary epithelial cells (HMEC), and all ENCODE cis-regulatory elements. The peak-to-gene correlation loops show the correlation between *ANXA2* and Enh1. Gene expression of *ANXA2* in matched scRNA-seq cells is depicted to the right of the browser track. **** denote *p*-value < 2.22e-16 (Wilcoxon rank-sum test). D. Find individual motif occurrences (FIMO) predictions within the *ANXA2* cancer-specific Enh1 and the *ANXA2* promoter showing the predicted transcription factors with the highest expression in the 0-Epi.Tumor cluster from patient 2. The gene expression of each TF specifically within the cancer epithelial cells is shown to the right of the table. E. Browser track view of chromatin accessibility at the *PRDX4* locus across epithelial cancer cells (Epi. Tumor) and all other cell-type clusters. Predicted enhancers of *PRDX4* are highlighted in light purple. The cancer-specific peak, Enh2, is denoted in red. The peak track below the browser track denotes all scATAC-seq peaks from this study (Peaks), regulatory elements found in human mammary epithelial cells (HMEC), and all ENCODE cis-regulatory elements. The peak-to-gene correlation loops show the correlation between *PRDX4*, and the peaks linked to this gene. Gene expression of *PRDX4* in matched scRNA-seq cells is depicted to the right of the browser track. **** denote *p*-value < 2.22e-16 (Wilcoxon rank-sum test). F. Find individual motif occurrences (FIMO) predictions within the *PRDX4* cancer-specific Enh2 and the *PRDX4* promoter showing the predicted transcription factors with the highest expression in the 0-Epi.Tumor cluster from patient 2. The gene expression of each TF specifically within the cancer epithelial cells is shown to the right of the table.

Another salient example of a cancer-specific enhancer linked to a male breast cancer upregulated gene is PRDX4. We selected PRDX4 for this example because this gene is involved in breast cancer metastasis53,54. We found that PRDX4 is more highly expressed in MBC compared to FBC (log2FC > 1.0), and its expression in cancer-epithelial cells is significantly higher than in the non-malignant cell types (Fig. 4E). In this case, there are three enhancer peak-to-gene links to the *PRDX4* promoter (Fig. 4E). Among these three peak-to-gene links, enhancer 2 showed the highest enrichment in the cancer epithelial cells compared to the non-malignant cells (log2FC = 3.5 and FDR < 1e-5) and was also not present in normal mammary epithelial cells (Fig. 4E). Motif search revealed that *FOXP1, TFAP2A* and *RUNX1* as the most significantly enriched transcription factors in enhancers 1-3, respectively, and *ZKSCAN1* was the most enriched transcription factor at the promoter of PRDX4 (Fig. 4F). Finally, we confirmed the expression of these transcription factors specifically within cancer epithelial cells using the scRNA-seq data, giving confidence to our results.

As we were looking through the 61 MBC genes that are linked to cancer-specific enhancers, we noticed a few instances of clusters of neighboring enhancers that had abnormally high levels of chromatin accessibility in the cancer-epithelial cell types. These clusters of enhancers were highly active and highly interconnected, reminiscent of super-enhancer function ^55,56^. For example, *LAMB3* is very highly expressed in MBC epithelial tumor cells, with more than five peak-to-gene links associated with its promoter (Supplementary Fig. S3A). The cluster of enhancer peaks near chr1:209,444,222-209,546,602 were significantly correlated to each other and individually linked to the promoter of *LAMB3*, suggesting that this may be a super-enhancer (Supplementary Fig. S3A). *LAMB3* has been previously related with focal adhesion for cell migration in breast cancer^57^, and its higher expression is associated with worse overall survival (OS) and disease-free survival (DFS) in pancreatic ductal adenocarcinoma (PDAC)^58^. Therefore, we posit high expression of LAMB3 is unfavorable for MBC patients and its elevated expression is sustained by the super-enhancer. Interestingly, we found a second cluster of cancer-cell specific enhancers that are linked with the expression of *CD47* and overlap with a previously annotated super-enhancer in breast cancer^59^ (Supplementary Fig. S3B). As seen with the previous example (*LAMB3*), the cluster of enhancer peaks near chr3:107,999,565-108,001,642, were highly interconnected and linked to the promoter of *CD47* leading to high levels of *CD47* gene expression (Supplementary Fig. S3B). CD47 is a key mediator of immune evasion and epithelial to mesenchymal transition in breast cancer and its high expression is related to worse disease free survival^60^. Although we did not intentionally set out to find super-enhancers, these examples highlight the potential for finding ‘super-enhancers’ at single-cell resolution. These data begin to explain how differentially expressed genes in MBC are regulated by cancer-specific enhancers, highlighting their potential as novel targets for therapeutic intervention.

## DISCUSSION

The results described herein exemplify the utility of our matched scRNA-seq and scATAC-seq dataset for uncovering clinically relevant mechanisms of gene expression in male breast cancer. We recognize that our study is limited to two ER+ male breast cancer patients but note that MBC represents only 1% of all breast cancer diagnoses, making it difficult to procure these rare tumor types. Moreover, we had the additional requirement of collecting live tumor specimens, on the day of surgical resection, to ensure high quality single-cell datasets. Thus, our study represents the first *matched* scRNA-seq and scATAC-seq dataset of MBC and represents a foundation for determining the regulatory logic of MBC. This is important because the 5-year mortality of MBC is higher than that of FBC^4^ and not enough is known about MBC at the molecular level, especially at single-cell resolution.

While the anatomical listing of cell types within MBC tumors was not the initial goal of this project, our high-quality scRNA-seq dataset enabled the detection of various cell types, including macrophages, fibroblasts, endothelial cells, and epithelial cells. The malignant cell types were detected by inferring copy-number variation from the scRNA-seq data, and notably we found that the non-malignant cell types showed similar gene expression profiles across patients (suggesting a similar biological state) while the cancer-epithelial cells showed more patient specific transcriptomes, pointing to the heterogeneity of MBC. Comparison of the cancer-epithelial cell transcriptomes to the female cancer-epithelial cell transcriptomes revealed distinctive gene expression programs as evidenced by the discrete clustering of the male versus female epithelial cells (Fig. 2). The top five upregulated genes were *RPS4Y1, TNFRSF12A, ISG20, S100A6, LGALS3*, whereas the top five downregulated genes were *XIST, FOS, TFF3, PBX1*, and *HSPA1B. RPS4Y1* was not expressed female epithelial cells since it was located on the Y-chromosome, confirming the soundness of differential gene expression analysis. Interestingly, the other up-regulated genes are associated with features of cell proliferation (*TNFRSF12A* in human hepatocellular carcinoma^61^), tumor progression (*ISG20* in clear cell renal cell carcinoma^62^), and cell survival (*LGALS3* in breast cancer^63^). Collectively the pathway analysis of all upregulated genes shows enrichment in hallmark pathways such as ‘*MYC targets*’ and ‘*mTORC1 signaling*’ (Fig. 2). Indeed, the upregulation of the *mTOR* pathway in male breast cancer has been shown before using bulk microarray studies^23^ suggesting that MBC patients may benefit from *mTOR* inhibitors. Conversely, the downregulated genes were enriched in the ‘*Estrogen Response Late*’ hallmark pathway, suggesting that although the male epithelial cells are ER+, the estrogen driven gene expression pathways may be muted. Similar to FBC, endocrine therapy targeting ER is the mainstay of systemic therapy for ER+ MBC, and these results indicate that MBC may be inherently less estrogen responsive. No data currently exist prospectively determining the efficacy of endocrine therapy in MBC patients. The ongoing ETHAN prospective, randomized trial (https://clinicaltrials.gov/ct2/show/NCT05501704), will compare endocrine therapy regimens in male breast cancer patients and this will be important to establish the role of endocrine therapy for male patients. Additionally, this trial will provide samples for translational correlates of how MBC respond to endocrine therapy, and to test how expression of genes identified in our study correlate with response.

The chromatin accessibility profiles enabled the systematic detection of regulatory elements that drive the distinctive gene expression programs within MBC. We demonstrated that there is widespread rearrangement of the enhancer landscape in MBC and that cancer-epithelial cells acquire de novo enhancer elements that drive the expression of male-specific genes. Starting from a broad survey of cis-regulatory elements across all cell-types, we were able to identify cancer-cell specific enhancer-to-gene links that are not typically active in normal cell types (Fig. 3). Examples such as *ANXA2* and *PRDX4* show significant enrichment in the chromatin accessibility of the cancer-specific enhancers compared to the non-malignant cell types, resulting in marked increases in gene expression (Fig. 4). Previous studies in female breast cancer have shown that increased *ANXA2* gene expression is associated with drug resistance and tumor recurrence^64^. Since *ANXA2* expression is even higher in MBC, our enhancer-promoter interaction model may provide a way to understand the mechanism by which MBC becomes resistance to standard therapy. Similarly, *PRDX4* is overexpressed in several cancers such as gastric cancer^65^, lung cancer^66^, and breast cancer^54^. In addition, elevated *PRDX4* expression is associated with worse overall and disease-free survival in female breast cancer patients^54^. Thus, the heightened expression of *PRDX4* in males versus females may underlie observed worse outcome for male patients.

Perhaps more interestingly, we saw evidence for super-enhancer activity at single-cell resolution. Super-enhancers are defined as large clusters of neighboring enhancers that have an unusually high occupancy of interacting factors and are thought to act synergistically with each other to promote the expression of their target genes^67^. Super-enhancers have gained much attention because they are known to regulate key cell identity genes, and in cancer are known to drive oncogene expression^56,68^. Within our list of 61 upregulated male versus female breast cancer genes linked to cancer-specific enhancers, we identified at least two super-enhancers; one linked to *LAMB3* and the other to *CD47* (Supplemental Fig. 3). Notably, the super-enhancer linked to *CD47* has been annotated before in breast cancer ^59^ and we show evidence of the increased activity of the constituent enhancers specifically within the cancer-epithelial cell types (Supplemental Fig. 3). *CD47* is a transmembrane protein that belongs to the immunoglobulin superfamily that enables binding adhesion to the extracellular matrix. The encoded protein binds the ligand thrombospondin and plays a role in membrane signal transduction^60,69^. In breast cancer *CD47* is associated with epithelial-mesenchymal transition and poor DFS^60,69^. Therefore, the high expression of *CD47* in MBC may suggests a possibility of targeting *CD47* as a therapeutic strategy to overcome immune evasion^69^. Although we may be underpowered to comprehensively annotate super-enhancers using single-cell data, the detection of super-enhancers at single-cell resolution is an interesting concept and inspires further investigation. Taken together, this study annotates clinically relevant regulatory networks in male breast tumors at single-cell resolution, providing a useful resource that expands our current understanding of the gene expression programs that underlie the biology of MBC.

## METHODS

### Human patient tumor dissociation

Fresh, never frozen or fixed, tumors were collected by UNC’s Tissue Procurement Facility and immediately transported to the lab on ice in DMEM/F12 media + 1% penicillin/streptomycin after surgical resection or core biopsy. Tumor specimens were dissociated and prepped as described in our previous work^37^. Briefly, tumor specimens were minced with two sterile razor blades and incubated overnight on a 37°C stir plate at 180rpm in the following digestion media: DMEM/F12, 5% FBS, 15mM HEPES, 1x Glutamax, 1x Gentle Collagenase/Hyaluronidase (Stem Cell Technologies, 07919), 1% penicillin/streptomycin, and 0.48 µg/mL Hydrocortisone (Stem Cell Technologies, 74144). After overnight digestion, tumor cells were pelleted at 1200rpm for 5 min at room temperature and washed two times with cold PBS + 2% FBS and 10mM HEPES (PBS-HF).

Removal of red blood cells was done by treating cell pellets with cold ammonium chloride solution (Stem Cell Technologies, 07850) diluted with 1 part PBS-HF. To ensure complete dissociation of tumor cells and removal of free DNA in the cell suspension, cells were treated with 0.05% Trypsin-EDTA and 200 µL 1mg/mL DNase I for one minute, followed by trypsin inactivation with PBS-HF and centrifugation. Cells were then washed with PBS-HF, filtered through a 100µm cell strainer and a final centrifugation was done. Cell pellets were resuspended in DMEM/F12 + 5% FBS using a volume that was dependent on the size of the final cell pellet, then filtered through a 40µm cell strainer. We measured cell viabilities for two patient samples with the Countess II Automated Cell Counter (Thermo Fisher, AMQAX1000), cell viabilities were 90% for the Patient 1 sample and 88% for the Patient 2 sample.

### Single-cell sequencing library preparation

Immediately following dissociation, cells were diluted to 1200 cells/µL in preparation for scRNA-seq and scATAC-seq library generation as described in our previous work^37^. For scRNA-seq libraries, 10,000 cells were used for library preparation using the 10x Genomics Single Cell 3’ kits: Single Cell 3’ GEM, Library & Gel Bead Kit v3 (PN-1000075), Chromium Chip B Single Cell Kit (PN-10000153), and Chromium i7 Multiplex Kit (PN-120262) following the manufacturer’s protocol.

For scATAC-seq, nuclei isolation of 500,000 cells was carried out following the Nuclei Isolation for Single Cell ATAC Sequencing protocol from 10x Genomics using a four-minute lysis time. Next, 10,000 nuclei were used for library preparation using the 10x Genomics Single Cell ATAC Kits: Chromium Single Cell ATAC Library & Gel Bead Kit v1 (PN-1000110), Chromium Chip E Single Cell ATAC Kit (PN-1000082), and Chromium i7 Multiplex Kit N, Set A (PN-1000084) following the manufacturer’s protocol. scRNA-seq and scATAC-seq libraries were sequenced using 10x Genomics’ suggested sequencing parameters on an Illumina NextSeq 500 machine by UNC’s Translational Genomics Lab.

### Single-cell RNA-seq quantification and quality control (QC)

Cell Ranger from 10x Genomics was used to generate raw and filtered feature barcode matrices for each patient sample. A Seurat object was built from the filtered feature barcode matrix for each patient sample by using the Seurat R package^41,70^. Quality control (QC) and doublet removal were performed separately for each patient dataset to select for high quality cells. First, outlier cells were defined in each of the following metrics: number of UMI counts) (< 5,000), number of genes expressed (< 2,000) and percent mitochondrial read count (> 25%). Outlier cells according to these criteria were removed before doublet detection. Then, doublet removal was performed by DoubletFinder^71^. After QC and doublet removal for each patient dataset, we applied Seurat’s *merge()* function^40^ to combine the individual patient datasets, forming the male breast cancer cohort presented in this study.

### Single-cell RNA-seq normalization, feature selection and clustering

Seurat’s *NormalizeData()* function^40^, with the normalization method set to “LogNormalize”, was used to normalize gene expression matrices. Seurat’s *FindVariableFeatures()* function^40^, with the selection method set to “vst” and the number of top variable features set to 2,000, was used to perform feature selection. Seurat’s *ScaleData()* function^40^ was used to scale the expression values for the top 2,000 variably expressed genes in the dataset before carrying out principal component analysis (PCA). We chose to regress out the percentage of mitochondrial genes when using Seurat’s *ScaleData()* function^40^. The top 2,000 most variably expressed genes were summarized by PCA into 50 principal components (PCs). Cells were then visualized in a UMAP embedding with Seurat’s *RunUMAP()* function^40^ using 30 PCs. Next, the shared nearest-neighbor graph was constructed by Seurat’s *FindNeighbors()* function^40^ using 30 PCs and Louvain clustering was performed with Seurat’s *FindClusters()* function^40^ with a resolution of 0.8. scRNA-seq UMAP embeddings were plotted in R^70^ using ggplot2^72^.

### Inference of copy number variation (CNV) from single-cell RNA-seq

Within each patient sample, estimated copy number events for each cell cluster were derived using the R package inferCNV^42^. Immune cell and endothelial cell clusters were used as a normal background for inferCNV. Remaining cell clusters were specified in the inferCNV annotations file to infer CNVs at the level of these clusters. The standard inferCNV algorithm was invoked with *infercnv::run()* with the following parameters of (cutoff: 0.1, scale_data: FALSE, HMM: FALSE, and denoise: TRUE). Epithelial cells were classified into epithelial tumor, epithelial unassigned, epithelial normal after plotting scatter plots with CNV values and correlation with top 5% cells with high CNV values^73,74^. Among nine clusters, two clusters were assigned as epithelial unassigned, and four clusters were assigned as epithelial tumor (Supplementary Fig. S2B).

### Single-Cell RNA-seq cell type annotation

The cell types were annotated with the R package SingleR^75^ based on reference transcriptomic datasets of pure cell types and gene signature enrichment obtained from Seurat’s *AddModuleScore()* function^40^. The normalized expression values built from human bulk RNA-seq generated and supplied by Blueprint and ENCODE were used as a reference dataset for SingleR^75^, which is available via R package celldex^75^. We built a MBC patient cohort dataset after merging the raw count matrices of both patients into a Seurat object by using Seurat’s *merge()* function^40^, we applied Harmony batch correction^38^, and finally clustering cells based on Louvain algorithm (described above). The resulting cell-type clusters in our merged dataset were assigned a cell type labeled based on the majority cell type in each cluster.

Among nine clusters in the full cohort of two MBC patients, six clusters were assigned as epithelial clusters since the most frequent cells of these clusters were epithelial cells. The other three clusters were a fibroblast cluster, an endothelial cluster, and a macrophage cluster. The mean number of RNA counts across all nine clusters was larger than 5,000 (Supplementary Fig. S1C) since outlier cells with UMI counts < 5,000 were removed at the QC step. Similarly, the mean number of features across for nine clusters was greater than 2,000 (Supplementary Fig. S1D) since outlier cells with features < 2,000 were also removed at the QC step.

### Differentially expressed genes in single-cell RNA-seq between female BC and male BC

ScRNA-seq data for female ER+ breast cancer data were downloaded from GSE161529^43^. The differentially expressed genes in single cells between 16 female ER+ breast cancer patients and two ER+ male breast cancer patients were identified by Seurat’s *FindMarkers()* function^40^ with parameters (logfc.threshold; 0.25, min.pct: 0.5, min.diff.pct: 0.25, and max.cells.per.ident: 500). Then markers were selected by an additional filter of Padj < 0.01. The heatmap of Fig. 2B includes top 25 upregulated genes in male BC and top 5 downregulated genes with additional criterion of Padj < 1e-12 to select smaller number of genes for visualization. All of the differentially regulated genes are tabulated in Supplemental Table 1. The enrichment of cancer hallmark gene sets^45^ were identified by hypergeometric tests with clusterProfiler enricher()^76^ with q-value ≤ 0.01.

### Single-cell ATAC-seq quality control (QC)

Cell Ranger ATAC from 10x Genomics was used to demultiplex raw base call files into FASTQ files and generate a filtered peak-barcode matrix containing detected cellular barcodes and a fragments file as in the BED format for each patient sample. These fragment lists were read into the R package ArchR^39^ to perform quality control and doublet removal. To enrich for cellular barcodes, a threshold for log10(TSS enrichement+1) was set manually to 0.9 for both scATAC-seq samples while a sample-specific threshold of log10(number of unique fragments) was estimated using a Gaussian Mixture Model (GMM) for each scATAC-seq sample, as implemented in the R package mclust^77^. Barcodes below these thresholds in any of these metrics were excluded before doublet detection step. ArchR’s *addDoubletScores()* function^39^, with the knnMethod parameter of “UMAP”, was used to estimate doublet enrichment scores, and ArchR’s *filterDoublets()* function^39^, with the filterRatio parameter of 1.0, was used to filter out cellular barcodes as doublets.

### Single-cell ATAC-seq quantification, feature selection and integration with single-cell RNA-seq

An initial tile matrix of 500 bp genomic tiles across all cells was generated by the ArchR package^39^. We used the iterative latent semantic indexing^35,36,78^ (LSI) procedure implemented in the ArchR R package to reduce dimensions of the genomic tile matrix using two iterations with 25,000 variable features. 30 LSI dimensions were used to create a UMAP embedding with ArchR’s *addUMAP()*^39^ with the reduced dimension object obtained by Harmony batch correction^38,39^. By applying a corCutOff parameter of 0.75 to ArchR’s *additerativeLSI*(), we excluded LSI dimensions that have a correlation to sequencing depth greater than 0.75. scATAC-seq UMAP embeddings were plotted in R^70^ using ggplot2^72^.

ArchR’s *addGeneScoreMatrix()* function was used to estimate gene activity scores by considering accessibility within entire gene body and the activity of putative distal regulatory elements^39^. Seurat’s CCA implementation^41^ was executed via ArchR’s *addGeneIntegrationMatrix()* function^39^ to integrate scATAC cells with a scRNA cells by assigning each of the scATAC-seq cells a cell-type cluster identity from the matching scRNA-seq data, an associated label prediction score, and an imputed transcriptome. Cells with label prediction scores less than or equal to 0.5 were exclude before getting marker features for each cell-type cluster and calling peaks from pseudo-bulk replicates. For each inferred cell-type cluster, pseudo-bulk replicates were generated using the R package ArchR^39^ and pseudo-bulk peak calling was performed using MACS2^79,80^. Peak calls from each inferred cell-type cluster were merged into a universal peak set using ArchR’s default iterative overlap procedure. ArchR’s *plotBrowserTrack()* function^39^ was used to plot genomic browser tracks visualizing the chromatin accessibility across cell-type clusters, the peak locations of pseudo-bulk ATAC-seq peaks, and peak-to-gene correlations.

We applied additional filtering step to select highly confident epithelial tumor clusters for the downstream analyses including peak-to-gene correlation analysis, identifying cancer specific distal peaks, and predicting transcription factor occupancy at select putative enhancer regions. Two clusters assigned as epithelial unassigned (i.e., 1-Epi. Unassigned and 8-Epi. Unassigned) were removed since these clusters could be a mixture of tumor cells and normal cells. Note that the number of cells in the first epithelial unassigned cluster (i.e., 1-Epi. Unassigned) was around 500 cells and the number of cells in the second epithelial unassigned cluster (i.e., 8-Epi. Unassigned) was less than 500 cells. We did not use “the number of cells <500” as a filtering criterion, but these two epithelial unassigned clusters might have lower quality in peak calling. One cluster assigned as epithelial tumor (i.e., 6-Epi. Tumor) was removed since the mean number of ATAC fragments was less than 5,000 (Supplementary Fig. S1G). Note that this cluster had the lowest number of cells (Supplementary Fig. S1F).

### Peak-to-gene correlation analysis

ArchR’s *addPeak2GeneLinks()* function ^39^, with reducedDims set to “IterativeLSI” and dimsToUse set to “1:30”, was used to carry out the peak-to-gene correlation analysis to identify putative regulatory elements by correlating peak accessibility with imputed gene expression in scATAC-seq cells ^37^. To circumvent the sparsity of scATAC-seq data, low-overlapping aggregates of scATAC-seq cells were generated via a k-nearest neighbor procedure in the LSI space to ensure robust peak-to-gene associations and reduce noise.

Peak-to-gene links were selected by distal peak location and selection criteria of correlation ≥ 0.45 and FDR ≤ 1e-12 with the control of Benjamini-Hochberg (BH) method^81^. The number of remaining distal peaks was 11,719, participating in a total of 22,869 peak-to-gene links. The distal peak-to-gene links were clustered using k-means before being visualized in a heatmap using ArchR’s *plotPeak2GeneHeatmap()* function^67^.

The mean number of linked genes per distal peak and the total number of genes linked with distal peaks (Fig. 3E) was computed from a peak-to-gene metadata table having peak names defined by genomic coordinates, and corresponding gene names. The distribution of the number of linked genes per distal peak was used to compute p-value with Wilcoxon test to show that the numbers of linked genes per peak in epithelial cells and non-epithelial genes are statistically significant.

### Identifying cancer-specific distal peaks

To identify putative cancer-specific distal peaks as demonstrated in Fig. 3A and Fig. 3D, we used a genomic internal overlap analysis as previously described ^37^. The genomic coordinates of the distal peaks participating in the cancer-enriched peak-to-gene links were overlapped with a set of normal active enhancer marker H3K27ac peaks in human mammary epithelial cells (HMECs) obtained from ENCODE^49^. To find any overlaps including 1 bp overlap between the cancer-enriched peaks and the normal peaks in HMECs, we used the function *findOverlapsOfPeaks()* from the ChIPpeakAnno R package^82^ with the minoverlap parameter of 1. The cancer-specific peaks were defined as cancer-enriched peaks that did not overlap with any of the normal peaks in HMECs.

The cancer-specific distal peaks linked with upregulated genes in MBC compared with FBC were determined by upregulation criteria (log2FC > 1.0 and adjusted P-value < 0.01). Total of 61 cancer-enhancer link genes were selected (Fig. 4A). The gene expression of these 61 genes in FBC and MBC cells were visualized in heatmap form (Fig. 4B).

The normal peaks in HMECs and the full list of the candidate cis-Regulatory Elements (cCREs)^49^ derived from ENCODE data in hg38 were used for visualization of peaks around genes of interest in Fig. 4C and Fig. 4E. The gene expression values (log normalized count) of *ANXA2* and *PRDX4* in two groups (epithelial tumor clusters vs. non-epithelial clusters) were compared by the Wilcoxon test.

Motif enrichment analysis was performed between cancer-specific distal enhancer peaks and other distal enhancer peaks with ArchR’s *addMotifAnnotation()* function^39^ using CisBP (Catalog of Inferred Sequence Binding Preferences)^83^ and *peakAnnoEnrichment()* function^39^ with the cutOff of “FDR < 0.01 & log2FC > 1.0”. The top 10 motifs enriched in cancer-specific distal enhancer peaks included *FOXA1* and *TFAP2C* (Supplementary Table S3).

### Predicting transcription factor occupancy at select putative enhancer regions

The genomic location of the enhancer of *ANXA2* was chr15:60,223,133-60,223,633 (Enh), whereas the genomic locations of three enhancer of *PRDX4* were chrX:23,434,983-23,435,483 (Enh1), chrX:23,525,503-23,526,003 (Enh2), chrX:23,822,316-23,822,816 (Enh3).

Bedtools^84^ *getfasta()* was used to extract the sequences of the select putative enhancers in the malignant fraction of Patient 2, as shown in Fig. 4D and Fig. 4F, after accounting for single-nucleotide variants relative to the hg38 reference genome. To include single-nucleotide variants from the malignant fraction in our analysis, we used bcftools^85^ *mpileup* followed by bcftools^85^ *consensus* with a bam file containing fragments only from cellular barcodes present in in the Patient 2 malignant fraction. Cell Ranger’s *bamslice* was used to subset a position-sorted BAM file to make a malignant-specific BAM file for each patient sample. We ran Find Individual Motif Occurrences (FIMO)^86^ motif scanning with the putative enhancer sequences as input and with JASPAR2020 CORE^87^ containing curated and non-redundant transcription factor binding motifs for vertebrates. The statistically significant motifs with a *q*-value < 0.01 were sorted by the expression values of corresponding transcription factors in the malignant cells of Patient 2. The expression value of a transcription factor was calculated by the sum of log-normalized counts of the transcription factors across malignant cells in scRNA-seq, which was visualized in R^70^ using ggplot2^72^.

## Data availability

Raw data (10x FASTQs) and processed data for single-cell RNA-seq data and single-cell ATAC-seq have been deposited at the National Institutes of Health (NIH) Database of Genotypes and Phenotypes (dbGaP) (https://www.ncbi.nlm.nih.gov/gap/) and is available under the accession number phs003006.v1.p1.

## Code availability

All original code has been deposited on the Github, and it is publicly available at the Github repository maleBC_single_cells (https://github.com/hyunsoo77/maleBC_single_cells). Any additional information regarding data analysis is available from the lead contact.

## Acknowledgements

We thank the patients and their families. We thank the University of North Carolina (UNC) Tissue Procurement Facility for helping acquire tumor specimens, and the UNC Translational Genomics Lab for providing experimental support and helping us sequence libraries. We thank Michele Hayward at the Office of Genomics Research for helping us navigate the intuitional review board (IRB) process and the data submission process. This work was supported by grants from the NIH/National Cancer Institute (R01-CA273444-01), the Susan G. Komen Breast Cancer Foundation (CCR19608601), and the Department of Defense CDMRP Breast Cancer Research Program (BC180450) to H.L.F.

## Author Contributions

H.L.F, K.W, and P.M.S. conceived and the study.K.W and P.M.S. carried out tumor sample procurement and generated the scRNA-seq and scATAC-seq libraries. H.K designed and performed the computational analysis with input from H.L.F. and M.J.R. The manuscript was written by H.K, K.W, M.J.R, and H.L.F with input from all authors.

---

**Supplementary Figure S1.**
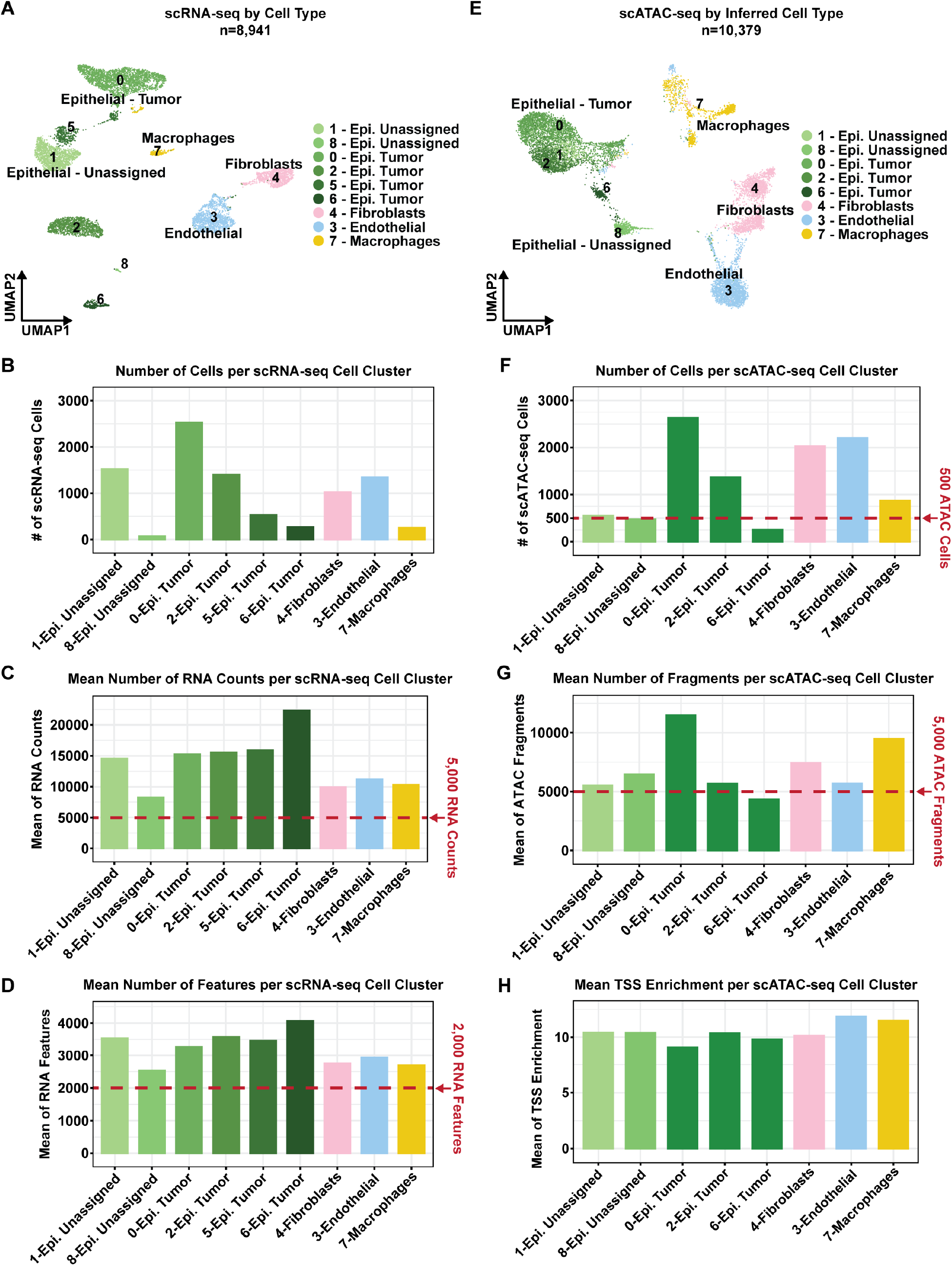
Quality metrics for scRNA-seq and scATAC-seq. A. scRNA-seq UMAP plot colored by cell-type, as in Fig. 1B, and numbered by cell-type cluster. B. Bar chart showing the number of scRNA-seq cells in each cell-type cluster. C. Bar chart showing the mean number of RNA counts per cell-type cluster. Red line denotes the cutoff threshold of 5,000 RNA counts. D. Bar chart showing the mean number of RNA features per cell-type cluster. Red line denotes cutoff threshold of 2,000 RNA features. E. scATAC-seq UMAP plot colored by cell type and numbered by cell-type cluster prior to filtering out low quality cells. Three cell-type clusters 1-Epi. Unassigned, 8-Epi. Unassigned, and 6-Epi. Tumor were filtered out in the UMAP in Fig. 1B. F. Bar chart showing the number of scATAC-seq cells in each cell-type cluster. The red line denotes the cutoff threshold of 500 ATAC cells. G. Bar chart showing the mean number of fragments per scATAC-seq cell-type cluster. Red line denotes the cutoff threshold of 5,000 ATAC fragments. H. Bar chart showing the mean TSS enrichment per scATAC-seq cell-type cluster.

**Supplementary Figure S2.**
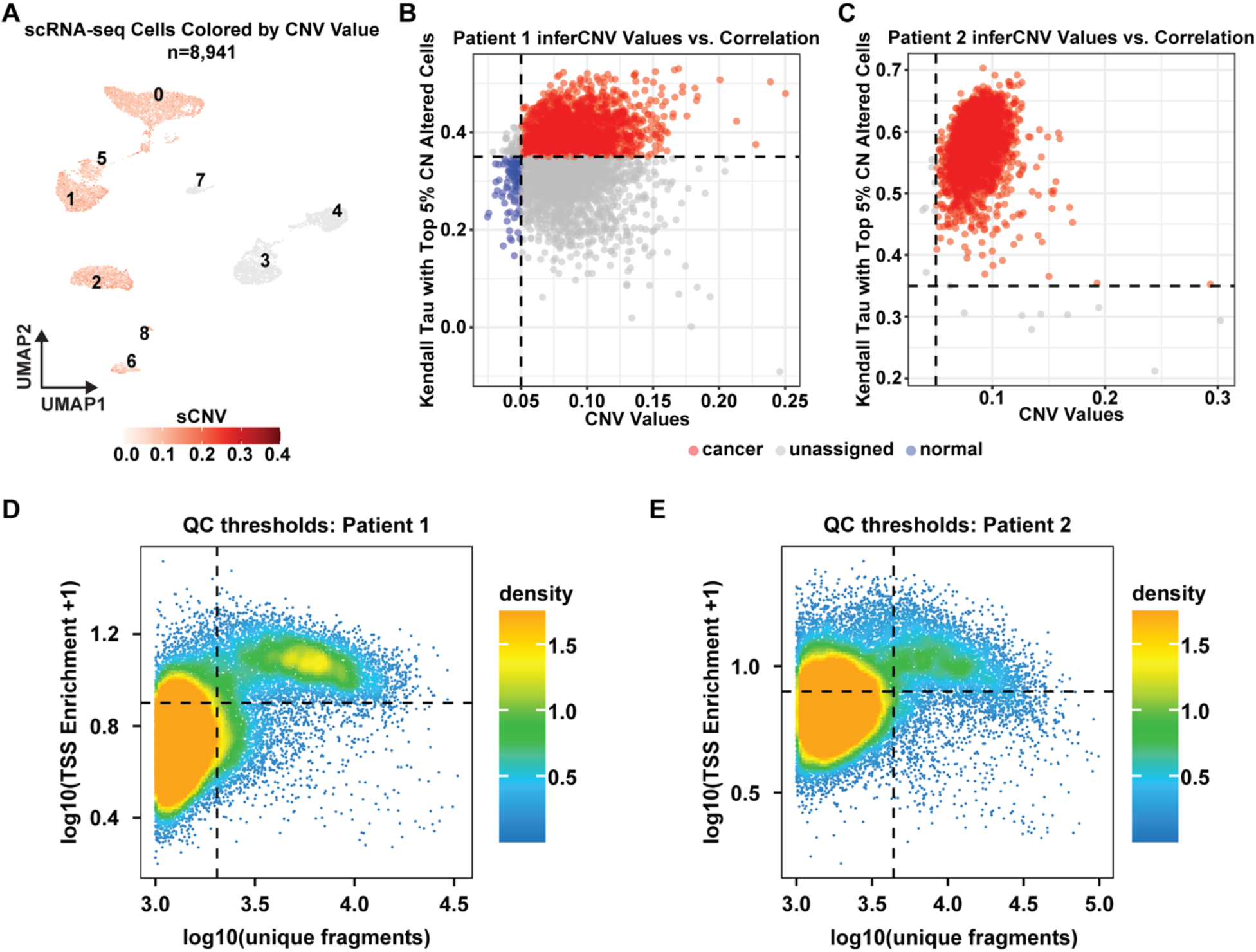
CNV Thresholds for scRNA-seq Cells and QC Thresholds for scATAC-seq Cells. A. scRNA-seq UMAP plot colored by CNV value. B. Scatterplot of Patient 1 inferCNV values vs. correlation. Correlation is measured as Kendall Tau with top 5% CN altered cells. Cells in red are considered cancer cells, cells in grey are considered unassigned cells, and cells in blue are considered normal cells. C. Scatterplot of Patient 2 inferCNV values vs. correlation. Correlation is measured as Kendall Tau with top 5% CN altered cells. Cells in red are considered cancer cells, cells in grey are considered unassigned cells, and cells in blue are considered normal cells. D. Scatterplot of barcode QC for Patient 1 scATAC-seq based on thresholds of TSS enrichment and log10(unique fragments). E. Scatterplot of barcode QC for Patient 2 scATAC-seq based on thresholds of TSS enrichment and log10(unique fragments).

**Supplementary Figure S3.**
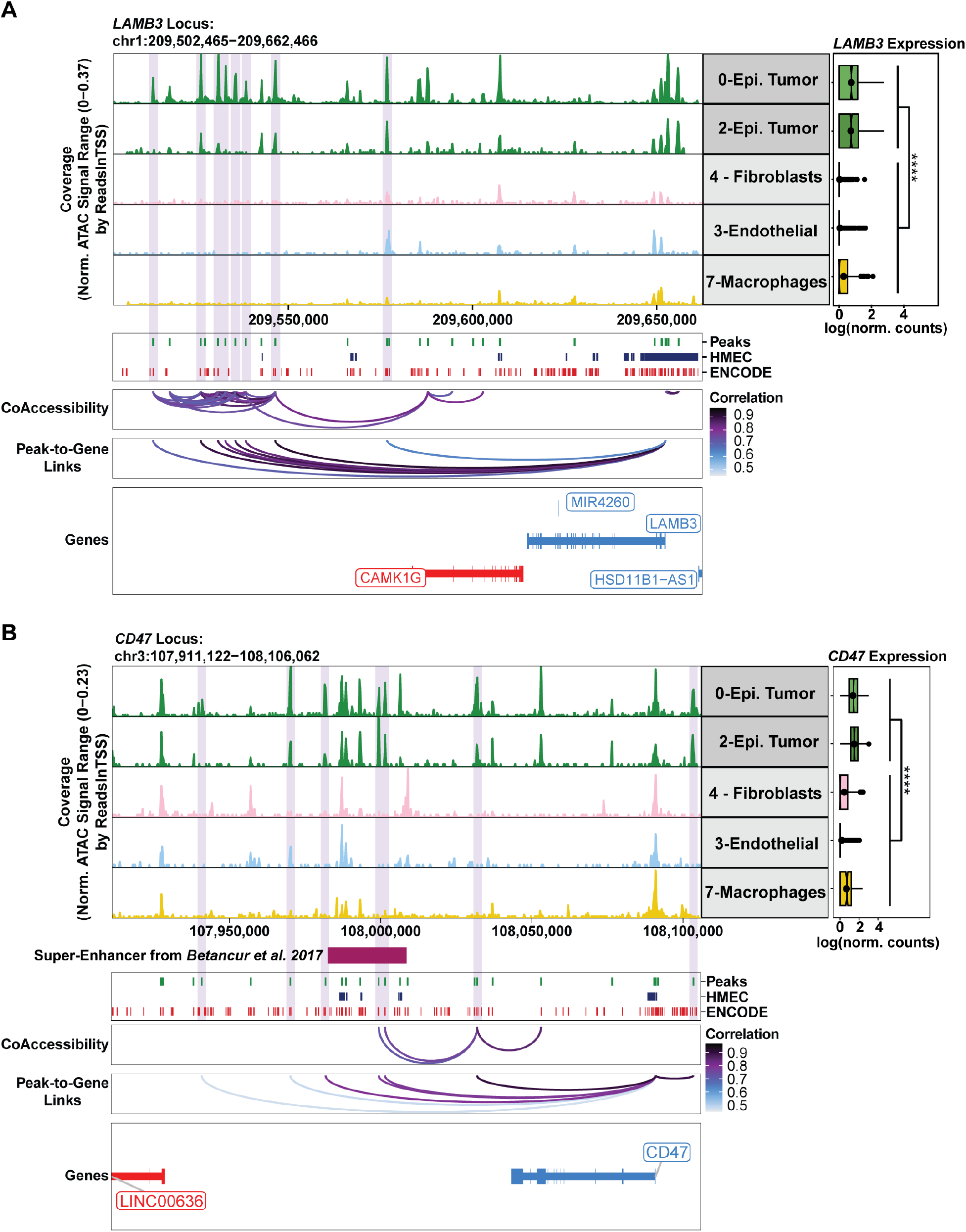
Peak-to-Gene Link Analysis Identifies Potential *LAMB3* Super-Enhancer and Confirms Previously Annotated *CD47* Super-Enhancer. A. Browser track view of chromatin accessibility at the *LAMB3* locus across epithelial cancer cells (Epi. Tumor) and all other cell-type clusters. Predicted enhancers of *LAMB3* are highlighted in light purple. The peak track below the browser track denotes all scATAC-seq peaks from this study (Peaks), regulatory elements found in human mammary epithelial cells (HMEC), and all ENCODE cis-regulatory elements. The peak-to-gene correlation loops show the correlation between *LAMB3* and the peaks linked to this gene. Gene expression of *LAMB3* in matched scRNA-seq cells is depicted to the right of the browser track. **** denote *p*-value < 2.22e-16 (Wilcoxon rank-sum test). B. Browser track view of chromatin accessibility at the *CD47* locus across epithelial cancer cells (Epi. Tumor) and all other cell-type clusters. Predicted enhancers of *CD47* are highlighted in light purple. The *CD47* super-enhancer from Betancur *et al*.^59^ is denoted in magenta. The peak track below the browser track denotes all scATAC-seq peaks from this study (Peaks), regulatory elements found in human mammary epithelial cells (HMEC), and all ENCODE cis-regulatory elements. The peak-to-gene correlation loops show the correlation between *CD47*, and the peaks linked to this gene. Gene expression of *CD47* in matched scRNA-seq cells is depicted to the right of the browser track. **** denote *p*-value < 2.22e-16 (Wilcoxon rank-sum test).

